# The effect of habitat type on population distribution and abundance of Rothschild’s Giraffe (*Giraffa camelopardalis rothschildi*) in Ruma National Park and Mwea National Reserve in Kenya

**DOI:** 10.1101/2021.04.30.442177

**Authors:** George N. Gathuku, David O. Chiawo, Charles M. Warui, Cecilia M. Gichuki, Innocent O. Ngare

## Abstract

The Rothschild’s giraffe is currently listed as vulnerable by the International Union for Conservation of Nature (IUCN). This is attributed to the loss of habitat due to human activities. This study examined the effect of habitat type on population structure and distribution of Rothschild’s giraffe in Ruma National Park (RNP) and Mwea National Reserve (MNR) in Kenya. The study employed road transects to collect data on the number, age class and sex distribution in three habitat types, open, medium and closed. Data was collected along three road transects of equal lengths measuring 14.2 km in each site (RNP and MNR) for comparison. A driving speed of 20 km per hour was maintained along each transect for standardization of survey effort and a maximum giraffe detection rate. Photographic capture of the coat patterns of the right side of all the giraffes sighted within 500 m from the transect was done for identification of age classes. The field visits were replicated 12 times for each transect giving 36 replications for each site spread equally through wet and dry seasons from March 2017 to November 2018. The effect of habitat type on population structure and distribution was analysed using ANOVA and Tukey HSD to test for significant differences. T-test was used to compare the mean population size of giraffe across the wet and dry seasons. Coat pattern analysis for age class identification was done using WildID software. The findings indicated that MNR had more males to females compared to RNP that registered more females and calves. Habitat type had a significant effect on the distribution of giraffes. The giraffe population showed a preference for medium habitat types. The findings are key for the management of habitat quality for giraffe populations at the interface where conservation areas overlap with human land use.

## INTRODUCTION

Africa serves as the cradle of the three giraffe sub-species as well as their origin. The observed decline in the giraffe population is consistent within the continent and across the world. Various giraffe subspecies are currently occurring in twenty one countries across Sub-Saharan Africa. These countries include Niger, down to Central, East and South Africa. This is made possible by their versatility in nature enabling them to survive and thrive in areas with minimal tree cover (1). According to (2) giraffes are extant in Cameroon, Chad, Botswana, Congo, Kenya, Ethiopia, Mozambique, Zambia, Zimbabwe, Uganda, and the Central Republic of Africa, Namibia, Niger, South Africa, Somalia, Tanzania, Angola and Sothern Sudan. Giraffes are believed to be extinct in many countries to include; Guinea, Mauritania, Eritrea, Senegal and Nigeria. The main threats facing the giraffe population have been sighted to be habitat loss and land fragmentation (3). East Africa and Kenya specifically remain home to three subspecies of giraffes; Maasai giraffe (*Giraffa camelopardalis tipplekirschi*), Rothschild’s giraffe (*Giraffa carmelopardalis rothschildi*) and Reticulated giraffe (*Giraffa carmelopardalis reticulate*) (4–6). Home ranges for giraffes often overlap and they vary across Africa and this is influenced by several environmental factors including, type of habitat and the amount of rainfall received, availability and quality of forage, herbivore verse vis predator density, as well as human influences including poaching and populations growth (7,8).

The Rothschild’s giraffe was first described by Sir Walter Rothschild in the 18^th^ century, by then, these giraffes were ranging feely and abundant across Kenya, Uganda and Sudan. In the late 1970s and early 1980s, Rothschild’s giraffes were relocated from Soi Ranch in the Rift Valley of Kenya to Ruma National Park, Nakuru National Park and later to Mwea National Reserve due to increasing human activities and settlements (9). According to (2), giraffe’s calves are usually preyed on by lions within the early years of their life, in a study conducted in Lake Nakuru National Park, and this impacts significantly on the population size of the Rothschild’s giraffe in this protected area. These accrued dynamic uncertainties have impacted the population of Rothschild’s giraffe by a 40% reduction from their existing previous population index. Currently, Rothschild’s giraffe is extinct in Sudan and there is only a small population remaining in Kenya and Uganda (10). The Rothschild’s giraffe is the second most endangered giraffe subspecies with less than 670 individuals surviving in the wild, of which 60% are in Kenya (11).

Rothschild’s giraffe population has continuously reduced in East Africa for the last thirty years (12). Since 1800, giraffe feeding and survival range have continued to shrink due to land fragmentation across Africa (2), hence giraffe decline by 40% in the last thirty years (13,14). This has been attributed to the high rate of human population increase which eventually needed areas to settle and cultivate for their basic livelihood needs (15). Due to the increasing need for agricultural settlement land and land conversion to privately owned cattle ranches, land fragmentation became so high, gradually increasing the giraffe’s vulnerability (5,16). By the year 2010, Rothschild’s giraffe population in the wild was less than 670 individuals (10). Over 500 of these giraffes were found in Kenya hence making the country an important stakeholder in the conservation and protection of the subspecies and its habitat (4). In June 2010, the IUCN declared the Rothschild’s giraffe subspecies as endangered (17).

Rothschild’s giraffe is historically known to have inhabited areas within eastern parts of Uganda and Southern parts of Sudan and some areas in the Rift valley in Kenya. However, this sub-species was eventually displaced from the original habitat due to human encroachment (2). Giraffes relate positively to arid and semi-arid habitats due to their browsing behaviour. However, research shows that their productivity is higher in humid environments rich in vegetative cover (18). Giraffes mostly feed on leaves, flowers, twigs and fruits, therefore, shaping the vegetation within the savannah ecosystems (19). Wooded grassland, which are rich ecosystems for their habitation are threatened by human anthropogenic factors that include forest fires, agriculture & habitat degradation (13).

In Kenya, Rothschild’s giraffes inhabited part of the Rift Valley on a land that was known as Soy Ranch or Soi Ranch which was an 18,000 acre Ranch before they were translocated to Ruma National Park and Lake Nakuru National Park in the late 1970s and early 1980s (20) for safety, due to human encroachment and settlement. Currently, in Kenya and Uganda, Rothschild’s giraffe numbers are estimated to be 1,671 individuals (14). Giraffes cover long distances in search of forage and mates, this has gradually increased the conflict between man and the giraffes as the demand for agricultural land increased and vast lands fragmented for infrastructure development and settlement, consequently reducing giraffe feeding range would negatively affect their breeding (21,22). Giraffes are megafaunas that browse on a variety of plants across their feeding ranges, mostly these ranges are usually replaced by crop farms which in turn reduce the variety of browse for the giraffes (19). In Kenya, they are faced with many extrinsic threat factors including; habitat loss, poaching, climate change, infrastructure development, inbreeding due to population isolation and intrinsic factors; loss of migration corridors, dietary complications and interspecific competition as well as diseases, that lead to reduced carrying capacity (23). Historically, Rothschild’s giraffes inhabited the Western parts of Kenya, but all known wild populations were extremely reduced mainly due to agricultural developments (14,23).

In Kenya, Rothschild’s giraffe populations were introduced in private and public areas, MNR and RNP in the late ‘70s and early ‘80s; RNP received 27 giraffes in total, 5 males and 22 females in 1983 (20) to rescue the giraffes from the human-induced threats (15). Despite being vulnerable and the ongoing threats, their population has been less studied and limited data exist to give clear indications of their abundance and distribution in Kenya (5). The study examined the effect of habitat types on population structure and distribution of Rothschild’s giraffe in Ruma National Park (RNP) and Mwea National Reserve (MNR) in Kenya. The findings are key for the management of habitat quality for giraffe populations at the habitat interface where conservation areas overlap with the fast-changing land use systems in human settlement areas.

## MATERIALS AND METHODS

### Site description

Data collection was conducted in Ruma National Park (RNP) 0°38’36’’S, 34°16’48”E; 1600 m ASL, located in South-Western Kenya, in Homabay County and Mwea National Reserve (MNR) 0°49’05”S, 37°37’19”E; 1100 m ASL, located in Embu County (Fig 1). RNP covers an area of 120Km^2^ with the majority of the Southern parts of the park dominated by wooded grasslands, with *Balanities aegyptica* (Egyptian balsam) and *Acacia drepanolobiam* (Whistling thorn) recording the highest numbers. Land use and the dominant human activities in the two areas is exclusively agriculture, with farmers growing a variety of crops e.g. maize, beans and keeping of domestic animals. MNR covers an area of 42Km^2^ and has diverse vegetation types, with some of the areas covered by wooded grasslands and bushlands with *Commiphora africana* (African Myrrh), *Acacia mellifera* (Black thorn) and *Combretum exalatum* (Combretum) dominating the *shrub layer. Acacia tortilis* (Umbrella thorn), *Grewia bicolor* (False brandy brush) *and Grewia villosa* (Mallow-leaved cross-berry) dominated the open grasslands (24).

**Fig 1:**
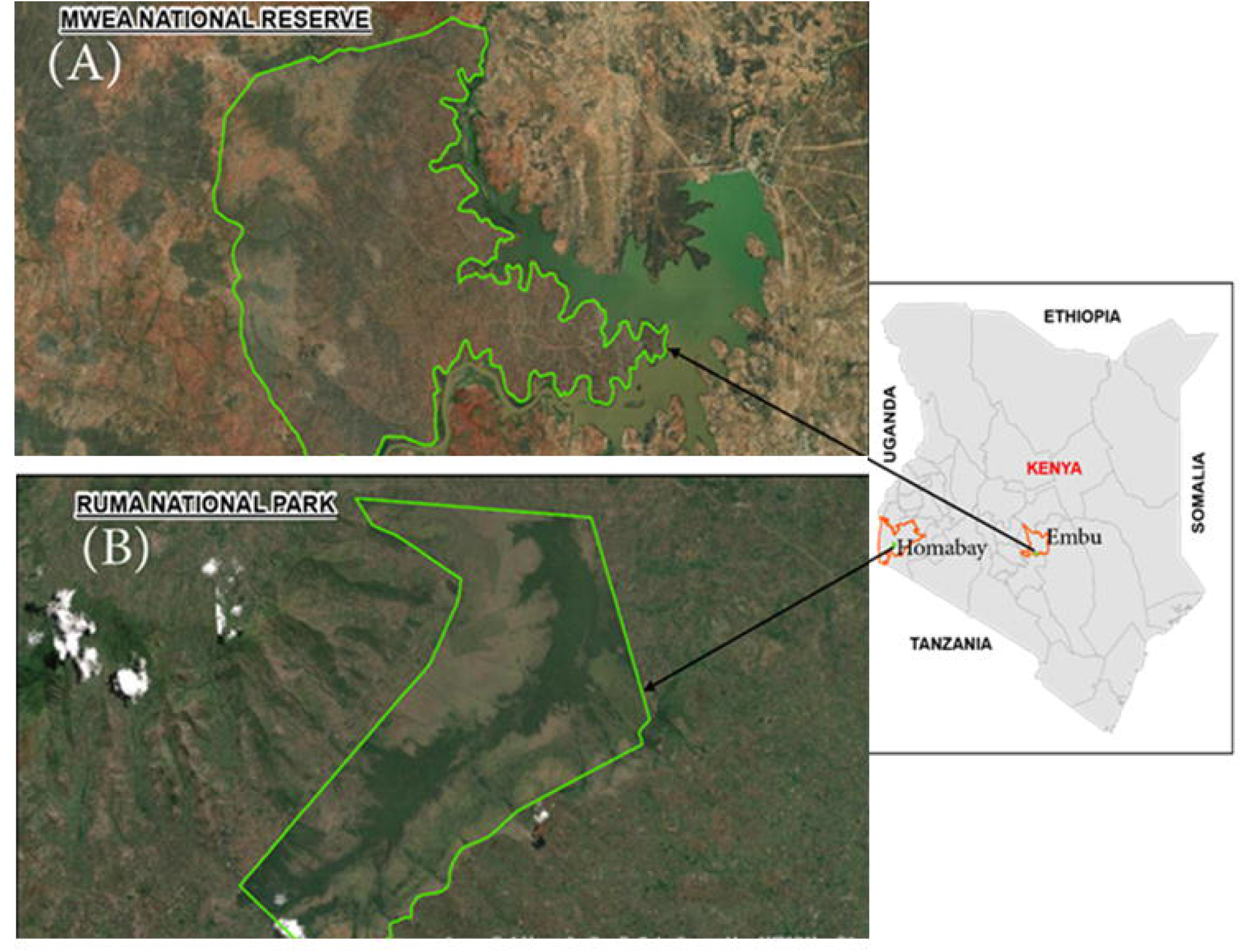
Map of the study areas, (A) Mwea National Reserve and (B) Ruma National Park, showing the boundaries and topography of the two protected areas during the period of study, March 2017 to November 2018.

### Data collection

The study employed 3 road transects of equal length measuring 14.2 km covering a total area of 41.2Km^2^ in each site (RNP and MNR) and photographic capture of the giraffe coat patterns. A driving speed of 20 kms per hour was maintained along each transect to ensure constant survey effort and maximum detection and encounter of giraffes (25). The driving speed was maintained at 20 Km/h along each transect for standardization of survey effort. Counts of giraffes sighted were made on both sides of the transects within 500 m range from each transect (26). On sighting the giraffes, numbers, habitat type, and sex were recorded. Photographic capture of the right side of the giraffe coat pattern was taken using a camera, Nikon Coolpix 900 for identification of the age class i.e., Adult, Sub-Adult and Calf. The images of the giraffe coat patterns were matched to available reference coat patterns of the age classes using WildID software (27). The field visits were replicated 12 times for each transect giving 36 replications for each site spread equally in wet and dry seasons for two annual years from March 2017 to November 2018. The survey pattern for the transects was alternated by selecting the transects randomly to reduce biases in day distribution.

### Data Analysis

The abundance of Rothschild’s giraffe in RNP and MNR was calculated as the cumulative number of individuals sighted along the transects. ANOVA and Tukey HSD post-hoc tests were used to test for significant differences. Captured images were run through the Wild ID software to identify the age classes of the individual giraffes recorded. Giraffe abundance was compared across age classes (adult, calf and sub-adults), and three habitat types (closed, medium and open) in RNP and MNR. T-test was used to compare abundance between dry and wet seasons and sex during the period of study. Comparison of wildlife populations across habitat types to determine any difference in distribution due to habitat type has also been applied by (28). Therefore, analysis was carried out separately for each site; RNP and MNR to allow for comparison in R statistical software (29).

## RESULTS

### Giraffe population structure and distribution

The total number recorded during the period of study in RNP were 314 giraffes, 221 (70.4%) adults, 72 (22.9%) sub-adults and 21 (6.7%) calves while in MNR they were 56 giraffes, 38 (67.9%) adults, 8 (14.3%) sub-adults and 10 (17.9%). The distribution in the number of calves varied between the two sites, (6.7% in RNP and 17.9% in MNR), RNP recorded a high number of females 181 (57.6%) while MNR recorded 24 (42.9%) females which was fewer than that of the males in the same protected area. MNR had a higher population of male giraffes (57.1%) as compared to the population of males in RNP (42%) relative to the female population (Table 1).

**Table 1:** Giraffe Total Numbers Distribution in Ruma NP and Mwea NR for data collected between March 2017 to November 2018, Ruma (N= 314), Mwea (N= 56).

| Category | Location |  |  |  | Total |  |
| --- | --- | --- | --- | --- | --- | --- |
|  | Ruma |  | Mwea |  |  |  |
|  | No. | % | No. | % | No. | % |
| Adult | 221 | 70.4 | 38 | 67.9 | 256 | 69.2 |
| Sub-adult | 72 | 22.9 | 8 | 14.3 | 80 | 21.6 |
| Calf | 21 | 6.7 | 10 | 17.9 | 31 | 10.2 |
| Male | 133 | 42.4 | 32 | 57.1 | 165 | 44.6 |
| Female | 181 | 57.6 | 24 | 42.9 | 205 | 55.4 |
| Total | 314 | 100.0 | 56 | 100.0 | 370 | 100.0 |

### The distribution of giraffe age classes and sex

There was a significant difference in the distribution of giraffes according to the age classes in MNR (F-2.862, N = 482, DF_2_ P<0.05), with a slightly higher number of Sub-Adults. While in RNP showed no significant difference in age class distribution (Fig 2). While more females were recorded in RNP, the number of females and males had no significant difference in both MNP and RNP.

**Fig 2:**
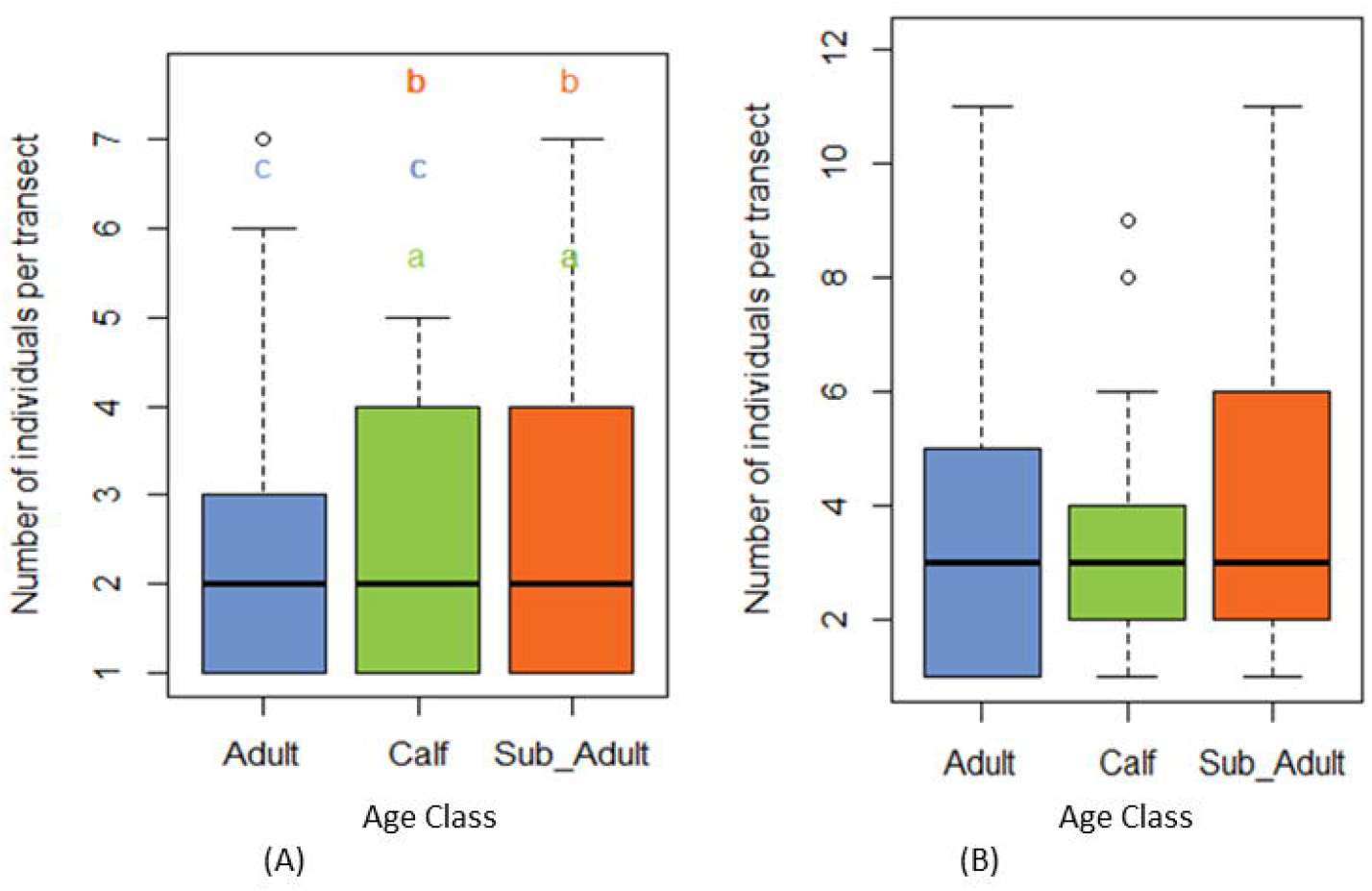
Boxplots comparing mean age classes of Rothschild’s giraffe distribution in Ruma National Park (A) and Mwea National Reserve (B) use types; Adult, Calf and Adult.

### Effect of habitat type on giraffe population distribution

Habitat type had a highly significant effect on the distribution of giraffes in Ruma, (F-106.2, N = 1723, DF_2_ P<0.001) and significant effect in MNR, (F-9.939, n=482, DF_2_ P<0.05). In RNP a significant difference was noted in all habitat types with medium habitat having high numbers followed by open then closed habitat. In MNR a significant difference was established between open and closed habitats and no difference between the pairs medium and closed, and open and medium (Fig 3).

**Fig 3:**
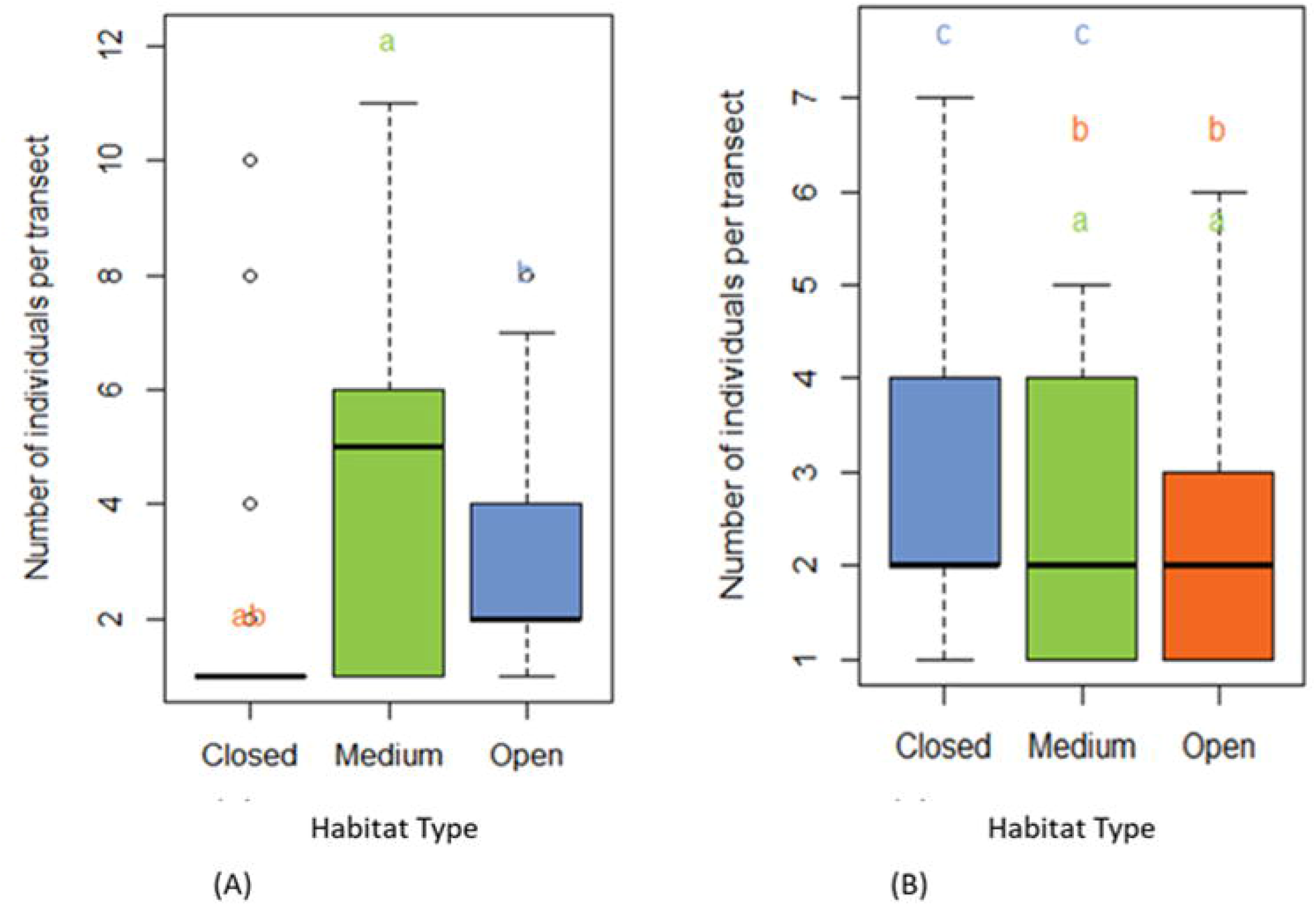
Boxplots comparing Rothschild’s giraffe distribution across habitat types in Ruma National Park (A) and Mwea National Reserve (B) Habitat types; Closed, Medium and Open.

### Effect of seasonality on giraffe population distribution

Seasonality showed a contrasting effect on giraffe distribution between MNR and RNP. MNR recorded a significantly high number during the wet season compared to the dry season, t= 2.6071, df = 324.12, P<0.05, while in RNP, a significantly high number was recorded during the dry season, t=14.178, df= 1723.5, P<0.001.

## DISCUSSION

### Giraffe population structure and distribution

The sex and age structure of giraffe populations differed significantly between RNP and MNR. MNR registered a very small number of calves, sub-adults and a large number of adult males. This could be attributed to predation and human encroachment. This agrees with (16) that established that direct risk such as predation significantly reduced the number of giraffes in Nakuru National Park and Soysambu conservancy in Kenya. The findings in MNR conforms to this indication, where the risk of encroachment from the community could be likely high and hence the low population of giraffes. The low number of calves and sub-adults in MNR could also be attributed to low resource availability. (30) and (2) indicated that a low number of calves in a giraffe population could be as a result of high competition due to low resource availability. Therefore, predation and community encroachment are likely contributing factors to the low numbers of calves in MNR. This consequently affects the number of sub-adults due to the small number that would reach their next life stage.

Giraffes are generally regarded as capital breeders and therefore they mostly rely on accumulated body reserve to satisfy the energy needs for reproduction as opposed to income breeders that usually meet their energy needs for reproduction in the short term (7). Ruma NP has enough forage for the giraffes all year round and the size of the park is adequate for the number of giraffes recorded hence a good indication for the high numbers of giraffes as opposed to MNR. Predation is one of the main causes of juvenile mortality in any given population in the wild (31,32), and this could be the case for MNR, however, predation is also a component of seasonal variation in juveniles and MNR is not an exception. It is expected that seasonal cases on reserve encroachment by the community are lower during the dry season, leopards being the main predators on giraffe calves in this Reserve. The presence of predators in any habitat would also result to a reduced reproduction success hence affecting the population growth of a species (14,17).

According to (33,34), in the wild giraffes exhibit social patterns that are characterised by changes in groups composition, where females are more in numbers than males being one aspect that makes a heard healthy for breeding (35), weak or lose inter-individual relationships and fewer preferential associations between giraffe age and sex classes have been previously reported (36). The finding of the high number of females in RNP agrees with the pattern explained, hence an indication of a healthy population and one ready for breeding. Habitat quality enhances adequate availability of browse in RNP leading to a high rate of breeding and survival of calves that translated into high numbers of sub-adults and consequently high numbers of adults (31). These species are likely to have sexual segregation, meaning that males and females use different habitats or forage and live in separate groups outside the mating season (37).

The five protected sites in Tanzania, including Lake Manyara National Park, Tarangire National Park, Manyara Ranch Conservancy, Lolkisale Game Controlled Area and Mtowambu Game Controlled Area, revealed a variation in adult female survival, which made the greatest contribution to variation in local giraffe population growth rates (31). This shares closely with the population structure observed in RNP where more females were recorded. Nonetheless, there is a biological trend that favour some female giraffes unlike the male giraffes in the lower altitudes compared to higher altitudes (38,39), hence the higher number of females in RNP than MNR. Therefore, the biological trend could be favouring the giraffe population in RNP, which is at the lower altitude

### Effect of habitat type on giraffe population distribution

In MNR, a high number of giraffes were recorded in medium habitats and low in closed habitats. This was an isolated case because MNR exemplifies more of the closed habitat within its ecosystem, hence no significant difference in the giraffe’s distribution in the Reserve. In RNP, giraffes preferred medium habitats, due to factors such as predation, competition and resource quality (40). Giraffe distribution in both medium and closed habitats was the same hence similar utilization of the habitats. This supports previous studies by (41,42) on ungulate species that showed that when temporal variation in adult survival is low, calf survival typically becomes the most important determinant of population growth rate. This behaviour is attributed to the fact that the males are comfortable across the habitats and they feel safe (43), while females prefer medium habitats where it is safer for the young ones from predators and quality browse and it’s availability for the young (40). Closed habitats recorded the least numbers of giraffes, these areas were dominated by males and very few females, calves and sub-adults. This concurs with (44), that giraffes inhabit dry, semi-arid and sub-tropical savannah environments that vary from open to closed habitats, but they avoid forest habitats. The reason for this would be predation and females keeping the calves away from threats or even competition from other species (45). Giraffes were recorded to invade farms and this heightened the human-wildlife conflict in both RNP and MNR communities, hence in reiteration the presence of human-oriented threats such as snaring and hunting being propagated towards these giraffes in both RNP and MNR (45).

### Effect of seasonality on giraffe population distribution

In RNP more giraffes were recorded in the three habitats during the dry season compared to the few numbers during the wet season. This could be attributed to the park floods during the wet season hence the giraffes stayed away from the low lying areas and instead remained on the higher grounds during the wet season resulting in few numbers recorded during this period (40). This is contrary to what was observed in MNR, with the reserve being well-drained, more giraffes were recorded during the rainy season as in the savannah woodland habitat as opposed to during the dry season. This could be due to forage availability or choice of forage during that particular period (46).

## CONCLUSION

The low giraffe population with an imbalance in population structure in Mwea National Reserved could be largely be attributed to risk factors relating to predation and human encroachment. Habitat type had a significant effect on the distribution of giraffes. The giraffe population showed a preference for medium habitat types characterised by mixed habitat structure including savannah and broken bushes.

## ACKNOWLEDGMENTS

Kenya Wildlife Service permitted the research in Ruma National Park and Mwea National Reserve. The Kenya National Council for Science and Technology authorized the study under permit number NACOSTI/P/19/23797/30241. The study acknowledges the role of the local community in Ruma and In Mwea who accepted to be interviewed. African Fund for Endangered Wildlife Kenya funded the study through Giraffe Conservation Foundation. Kenyatta University that approved this research for PhD studies, Murang’a University of Technology and Strathmore University for the staff time of the supervisors.

## Data sharing declaration

All the datasets used in this paper have been shared as occurrence datasets on the Global Biodiversity Information Facility portal (GBIF), doi.org/10.15468/97vcua for Ruma and doi.org/10.15468/dc34f2 for Mwea.

